# Crystal structure of the classical MHC-I molecule: insights into the MHC-I system in antiviral diseases in dogs

**DOI:** 10.1101/2021.01.04.425351

**Authors:** Yujiao Sun, Lizhen Ma, Shen Li, Yawen Wang, Ruiqi Xiao, Junqi Yang, Chun Xia

## Abstract

Only one classical MHC-I locus (aka DLA-88) evolved in dogs, and thus far, a total of 76 DLA-88 alleles can be divided into two categories. The first category consists of 60 alleles, and the second category consists of 16 alleles. The main difference between the two categories is the insertion of an amino acid in the α2 region of DLA-88 alleles. To elucidate the structure of the first category, in this study, the crystal structure of pDLA-88*001:01 was determined for the first time. The 3D structure and topological characteristics of the ABG of pDLA-88*001:01 with a CDV peptide were analyzed. The viral presentation profile and the binding motif of viruses presented by pDLA-88*001:01 were determined. Most importantly, there were no amino acid insertions in the α2 region of the first category, which changed the conformation of the D pocket and the docking of the TCR. The results suggest obvious differences between the two categories. Because of the variation in the α2 region, pDLA-88*001:01 showed distinctive features in the two categories. Due to the peptide-binding motif of pDLA-88*001:01, more than 320 high-affinity viral peptides were predicted from dog H7N9, CPV, CMV, CMV, and CDV strains. The results reveal that there are two kinds of structural MHC-I systems in dogs that are responsible for CTL immunity against viral diseases. The results provide knowledge for designing viral epitope vaccines in canines.

**Importance:** DLA plays an important role in the acquired immunity of organism. In previous study, the pMHC-I structure of dog was analyzed with DLA-88 self-peptide. In this study, we screened several viral peptides which can bind to DLA-88 and resolved the structure of the DLA-88 complex binding the CDV peptide. This study enriches the study of canine MHC-I molecular-presenting polypeptide-activated TCR, which is of great significance for the study of canine cellular immunity and anti-viral vaccine development.

## Introduction

Canines have a unique set of characteristics that meet human needs. Approximately fifty thousand years ago, humans domesticated wolves that evolved into today’s dogs (1). To date, approximately 450 breeds of dogs have been domesticated with a population of approximately 250 million (2). Dogs are the most successful animals to enter the human family and play various important roles in human society. Moreover, there are further increases in dog varieties, and the population of dogs is growing.

There is no doubt that the dog’s immune system is its defense against various viral diseases. Because dogs are mammals, although the genetic difference is significant, we can generally use the knowledge and concepts of human and mouse immunology to understand dog immunology (3). To date, the research and application of single and multiple vaccines for viral diseases, such as canine distemper virus (CDV), rabies virus, canine parvovirus (CPV), canine adenovirus (CAV) and canine coronavirus (CCV), have been completed successfully (4–8). However, because some pathogens do not have vaccines, and with the frequent occurrence of various events that affect vaccine immunization failure and the emergence of new infectious diseases (9), we need to have a better understanding of the immune system in dogs. These results will strengthen the prevention and control of dog viral diseases and prevent the cross-species transmission of important disease pathogens, such as the rabies virus, from dogs to humans and other animals.

Canine genome sequencing data show that dogs have a series of specific cellular immunity and antibody immune gene groups similar to those reported in mammals (10). In particular, the major histocompatibility complex class I (MHC-I)-T-cell receptor (TCR) recognition and T cell activation genes cluster with the antiviral immune response and the molecular MHC-I/II pathway can initiate a specific T cell immune response (11). Canine MHC (also known as dog leukocyte antigen, DLA) is located on chromosome 12 (12). There are three MHC-I loci on chromosome 12, namely, DLA-88, DLA-64 and DLA-12 (13), and another MHC-I locus on chromosome 18, namely, LA-79 (14). Among them, the three loci (DLA-64, DLA-12 and DLA-79) are currently considered to be nonclassical MHC-I genes (15,16); DLA-88 is the only classical MHC-I locus that presents antigen peptides, activates T cells, and completes cytotoxic T lymphocyte (CTL) immunity to kill virus-infected cells (17). Therefore, the MHC-I system in dogs is obviously different from that in other mammals (i.e., there is only one classical MHC-I locus (MHC-Ia, also known as DLA-88 in dogs)) (18). Polymorphisms of the DLA-88 locus have been investigated. In total, 76 dominant DLA-88 alleles were found in over 200 different breeds of dogs (19). According to the length of the antigen-binding groove (ABG) formed by DLA-88, that is, whether there is a single amino acid (Leu/His insertion) in the α2 region, 76 alleles can be divided into two categories. The first category consists of 60 alleles, and the second category consists of 16 alleles. The main difference between the two categories is the insertion of an amino acid in the α2 region of DLA-88. DLA-88*508:01 represents an allele with an insertion, and DLA-88*001:01 represents an allele without insertion. The DLA-88*508:01 molecule can bind to its self-peptide (20); the DLA-88*508:01 molecule has a preference of hydrophobic residues at positions 2 and 3 of the bound peptide, and peptides with positively charged residues at the C-terminus (21). Among them, two self-peptides, namely K11 (RFLDKDGFIDK) and K9 (KLFSGELTK), have a high binding affinity to DLA-88*508:01, and the structures pDLA-88*508:0l in complex with K11 or K9 have been solved (22). The results showed the amino acid anchoring mode of DLA-88*508:01 and suggested its potential TCR docking characteristics (22). Although there are still uncertainties about DLA-88, the study started the research on the dog MHC-I system from complementary DNA (cDNA) to protein. However, research on the antiviral and tumor DLA-88 protein levels has not been conducted in dogs.

Currently, the main factors threatening the health and survival of dogs are viral diseases, such as canine distemper. Since there are no specific therapeutic drugs for canine viral diseases, vaccinations are mainly used. The early inactivated vaccine of CDV has rarely been used because of its poor immunogenicity. In addition, the heterogeneous vaccine of measles virus cannot provide lasting immunity (23). Therefore, attenuated vaccines are mainly used to prevent canine distemper and other viral diseases. However, the attenuated vaccine also has some disadvantages, such as a weak virus returns as a strong virus and the interference by maternal antibodies during the first immunization, resulting in transient immunosuppression and encephalitis in immune animals. In recent years, the research and development of new vaccines, such as recombinant live virus vaccines, genetically engineered subunit peptide vaccines and nucleic acid vaccines, have started (24). The main research focus is to find a safe and effective vaccine, especially a new multi-epitope vaccine, to prevent canine distemper and other canine viral diseases.

To elucidate the structure of the first category of DLA-88, in this study, the crystal structure of pDLA-88*001:01 was determined. The topological characteristics of the ABG were analyzed. Most importantly, there is no amino acid insertion in the α2 region, which is involved in the changing conformation of the D pocket and in the docking of the TCR. Because of the variation in the α2 region, pDLA-88*001:01 also showed unique features. Finally, more than 320 high-affinity viral peptides were calculated from dog viral strains. The results reveal the structural characteristics of the MHC-I system in dogs and provide foundational knowledge for designing viral epitope-peptide vaccines.

## Results

### The unique DLA-88 alleles in dogs

The phylogenetic relationship among the classical DLA-88 alleles is shown in Fig. 1. DLA-88 alleles can be divided into 9 branches (labeled 1-9 from right to left). The homology of DLA-88 alleles is over 88%. Among them, there are amino acid changes in the α1 and α2 regions; in particular, there is an amino acid insertion in the α2 region. One kind of α2 region is 90 amino acids, and the other kind of α2 region is 91 amino acids due to an amino acid insertion (Fig. S1). There were 16 alleles with an amino acid (Leu or His) insertion, which were distributed in 1, 2, 3, 4 and 7 branches. The other 60 alleles α2 region were 90 amino acids, distributed in 2, 3, 4, 5, 6, 7, and 8 branches. The presence or absence of an amino acid insertion in the α2 region is an important feature of dog allelic polymorphisms.

**Fig. 1.**
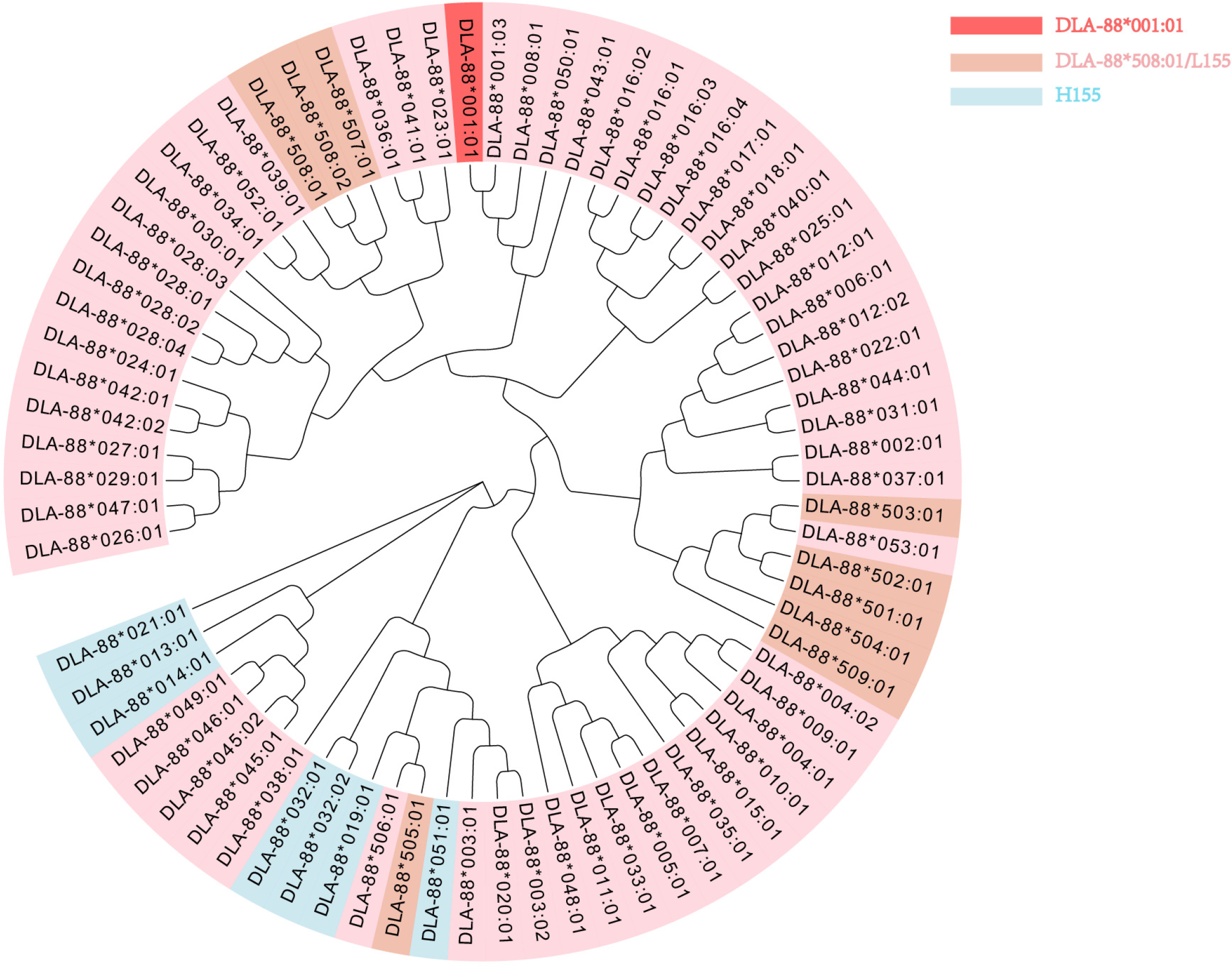
Phylogenetic relationship among the classical DLA-88 alleles. Nucleotide sequence-based phylogenetic tree of *DLA-88* alleles constructed by the neighbor-joining method. The α1 and α2 domains of 76 DLA-88 sequences that were released in IPD-MHC (https://www.ebi.ac.uk/ipd/mhc/alignment/help/) and NCBI were used to construct the phylogenetic tree. The DLA-88, DLA-79, DLA-64, and DLA-12 alleles are shown in pink, pale green, salmon and light blue background, respectively. The location of DLA-88*001:01 was confirmed with a crystal structure marked with a red background. The salmon color indicates that the 155th amino acid in the α2 region of the DLA-88 allele is Leu (Leu155) in. Light blue indicates that the 155th amino acid in the α2 region of the DLA-88 allele is His (His155).

### Overall structure of the pDLA-88*001:01 complex

Fourteen peptides, which could be presented by DLA-88*001:01, were synthesized for testing (Table S1). Among the predicted peptides, ten peptides could form peptide/DLA complexes (pDLA-88*001:01) by *in vitro* refolding. All of these ten peptides were used to screen the crystal structure of pDLA-88*001:01, and finally, the complex structure of DLA-88*001:01 with CDV_-RTI9_ was solved. X-ray data collection statistics are summarized in Table 1. Analysis of the crystal structure of the pDLA-88*001:01 complex showed two DLA-88*001:01 complexes with CDV-_RTI9_ in each asymmetric unit, termed DLA-88*001:01-CDV-_RTI9_ (aka pDLA-88*001:01). pDLA-88*001:01 contained the DLA-88*001:01 heavy chain (H chain, residues 1 to 276), dog β2m (residues 1 to 98) and CDV-_RTI9_ peptide (Fig. 2A). The DLA-88*001:01 H chain was composed of α1, α2, and α3 domains; the α1 and α2 domains form the ABG (Fig. 2B). The H chains of the α1 and α2 domains can be divided into two portions. One portion of these domains (α1, residues 51 to 55 and 58 to 86; α2, residues 139 to 151 and 153 to 176) forms helices located at the top of the ABG, and the remaining portion (residues 4 to 13, 22 to 29, 32 to 38, 95 to 104, 111 to 119, 122 to 127, and 134 to 136) forms a seven-stranded β-sheet at the bottom. Both the α3 domain (residues 186 to 193, 199 to 208, 214 to 219, 222 to 223, 229 to 230, 234 to 235, 241 to 248, 257 to 262, and 270 to 274) and β2m (residues 7 to 12, 22 to 31, 37 to 42, 55 to 57, 60 to 70, 78 to 83, 91 to 94) form two 7-stranded β sheets (Fig. 2). When DLA-88*001:01-CDV-_RTI9_ is compared with the already resolved structure of pDLA-88*508:01, the overall root-mean-square deviation (RMSD) is 0.521 Å (Fig. S2), and the RMSDs of the ABG, α3 domain and β2m are 0.468, 0.481 and 0.500 Å, respectively, indicating that there are consistent conformations in the two determined structures, but there are some important differences. For instance, there are a total of 18 hydrogen bonds between DLA-88*001:01 and DLA-β2m (Fig. 1C), which is greater than that (13 hydrogen bonds) in pDLA-88*508:01 (Fig. S3).

**Fig. 2.**
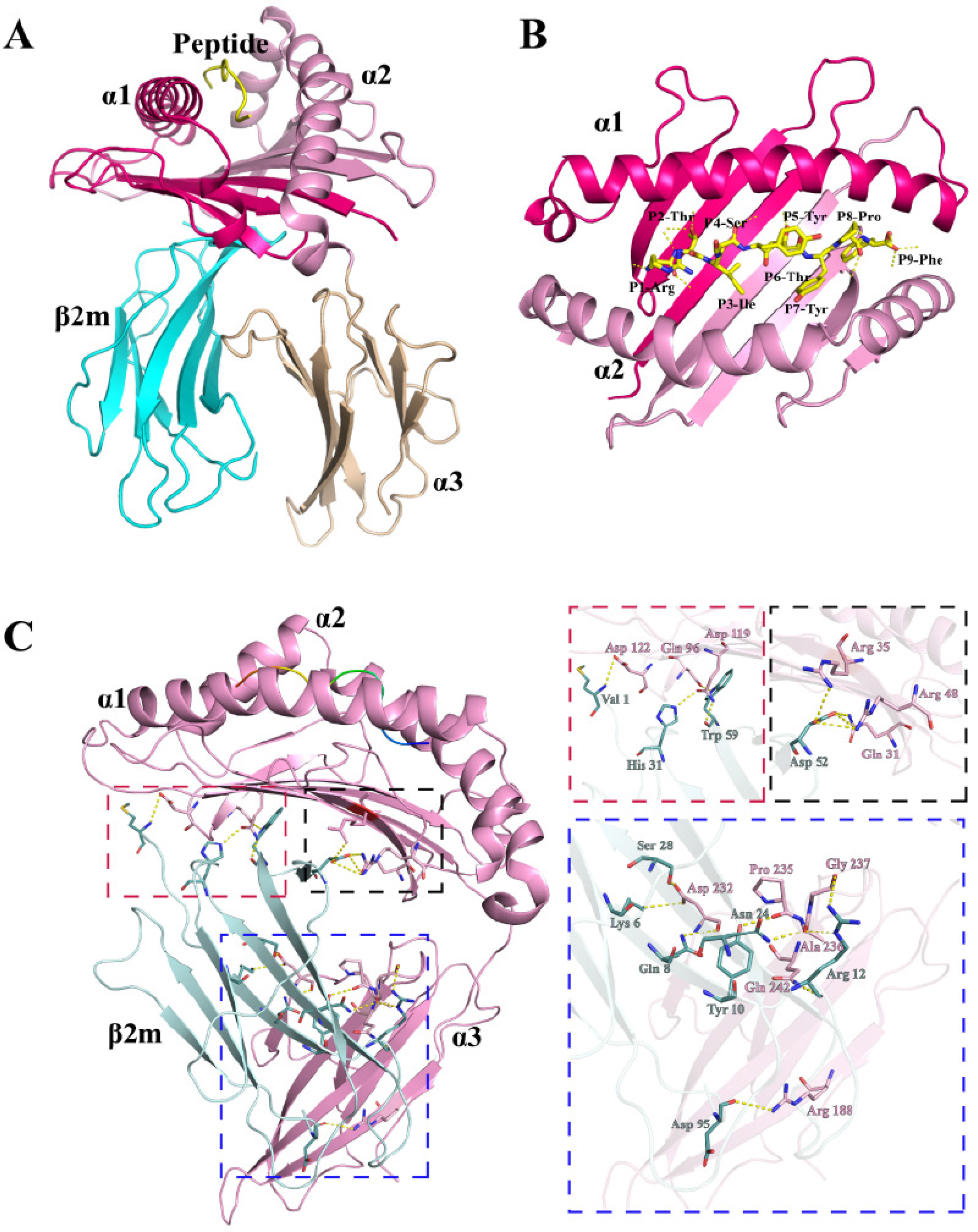
Structure of the domestic cat MHC-I molecule DLA-88*001:01. (**A**) Stereo view of the DLA-88*001:01 complex. The H chain, composed of the α1, α2, and α3 domains, is shown as a cartoon. The α1 domain is hot pink, α2 is light pink, and α3 is pink.; β2m is pale cyan. The CDV-H1_RF9_ (RTISYTYPF) peptide is shown as cyan stick models. (**B**) The overhead view of the α1 and α2 domains and CDV-H1_RF9_ (RTISYTYPF) peptides. (**C**) Interactions between the H and L chains of pDLA-88*001:01. The H chain is shown in pink, with the interactive residues shown as sticks and colored according to the atom type (blue, nitrogen; red, oxygen). The interactive residues on the L chain are white. The hydrogen bonds are shown as yellow dashed lines. All of the interactions and interactive residues are magnified within a corresponding color dotted box and labeled.

**Table 1.** Data collection and processing.

| Parameter or statistic | CDV-H1RF9 |
| --- | --- |
| Data processing |  |
| Space group | P2<sub>1</sub>2<sub>1</sub>2<sub>1</sub> |
| Cell parameters (Å) | a= 89.3, b= 93.0, c= 119.4 |
| Resolution range (Å) | 50.00-2.70 (2.75-2.70) |
| Total reflections | 152570 |
| Unique reflections | 50958 |
| Completeness (%) | 97.8 (96.9) |
| R<sub>merge</sub> (%)<sup>b</sup> | 0.201 (0.540) |
| I/σ | 7.308 (2.500) |
| R factor (%) | 25% |
| R<sub>free</sub> (%) | 28% |
| Bonds (Å) | 0.004 |
| Angles (°) | 0.926 |
| Average B factor | 17.013 |

$R_{\text{merge}} = \sum_{\text{hkl}} \sum_i |I_i(\text{hkl}) - \langle I(\text{hkl}) \rangle| / \sum_{\text{hkl}} \sum_i I_i(\text{hkl})$ , where $I_i(\text{hkl})$ is the observed intensity and $\langle I(\text{hkl}) \rangle$ is the average intensity from multiple measurements.
$R \text{ factor} = \Sigma (F_{\text{obs}} - F_{\text{calc}}) / \Sigma F_{\text{obs}}$ ; $R_{\text{free}}$ is the $R$ factor for a subset (5%) of reflections that was selected prior to refinement calculations and not included in the refinement.

### The length variation in the α2 domain found in DLA-88*001:01

To analyze the features of DLA-88*001:01, the amino acid sequences of DLA-88*001:01 and DLA-88*508:01 were aligned with typical peptide/MHC-I (pMHC-I) molecules from pig, human, cattle, rat, monkey, mouse and chicken (Fig. 3). Amino acid sequences of MHC-I alignment results show that the identities of DLA88*001:01 between mammalian MHC-I are more than 70%, and the identity of DLA88*001:01 with nonmammalian chicken pMHC-I is approximately 44%. According to the sequence alignment of the α1, α2 and α3 domains, the differences between the DLA-88*001:01 and DLA-88*508:01 alleles are mainly concentrated in the α2 domain. The most striking difference is an additional inserted Leu155 in DLA-88*508:01, which leads to the length variation in the α2 domains. The differences caused by this amino acid in the three-dimensional (3D) structures will be discussed in detail below.

**Fig. 3.**
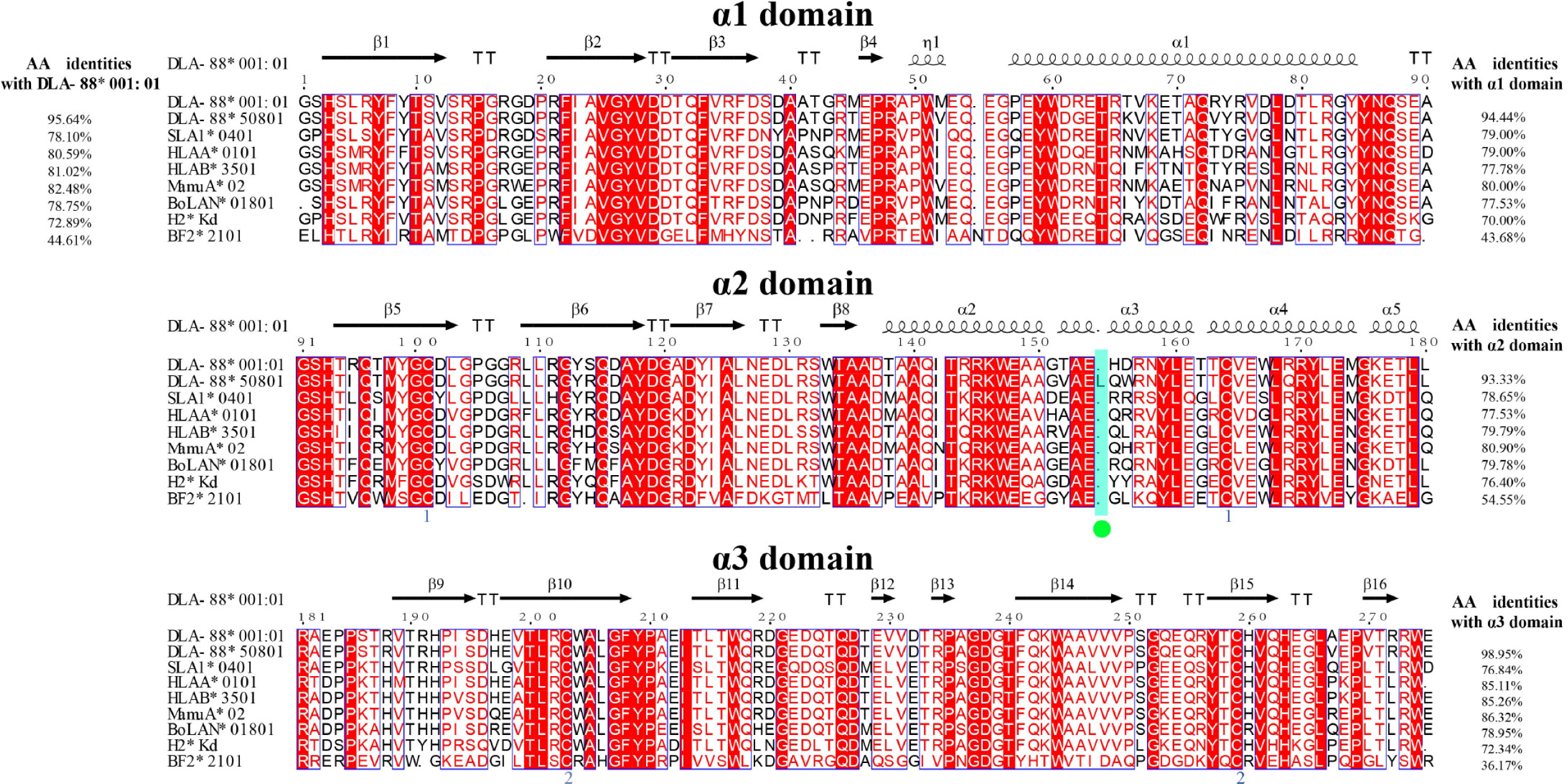
Structure-based amino acid sequence alignment of DAL-88*001:01 and other representative crystallized pMHC-I molecules, with the secondary structure elements indicated. The amino acid identities between DAL-88*001:01 and the listed MHC-I molecules are shown at the end of each sequence. The different residues of DAL-88*50801 with other sequences are shown with green circles below the sequence.

### The conformation of ABG in DLA-88*001:01

The extrinsic structure of pDLA-88*001:01 is similar to that of other resolved classical pMHC-I structures, and the ABG is divided into six pockets (pockets A-F, Fig. 4). The A-F pockets contain P1, P2, P6, P3, P7 and P9 amino acids, respectively, and the interactions between CDV-_RTI9_ and the ABG are listed in Table 2. The A pocket consists of eight amino acids, namely, Leu5, Tyr7, Tyr59, Glu63, Tyr159, Cys164, Trp167 and Tyr171, which fix the position of P1-Arg via hydrogen bonding and strong van der Waals (vdW) forces. The A pocket and CDV-_RTI9_ PN in pDLA-88*001:01 displayed strong interactions, such as the hydrogen bond network formed by Tyr7, Tyr159, and Tyr171.

**Fig. 4.**
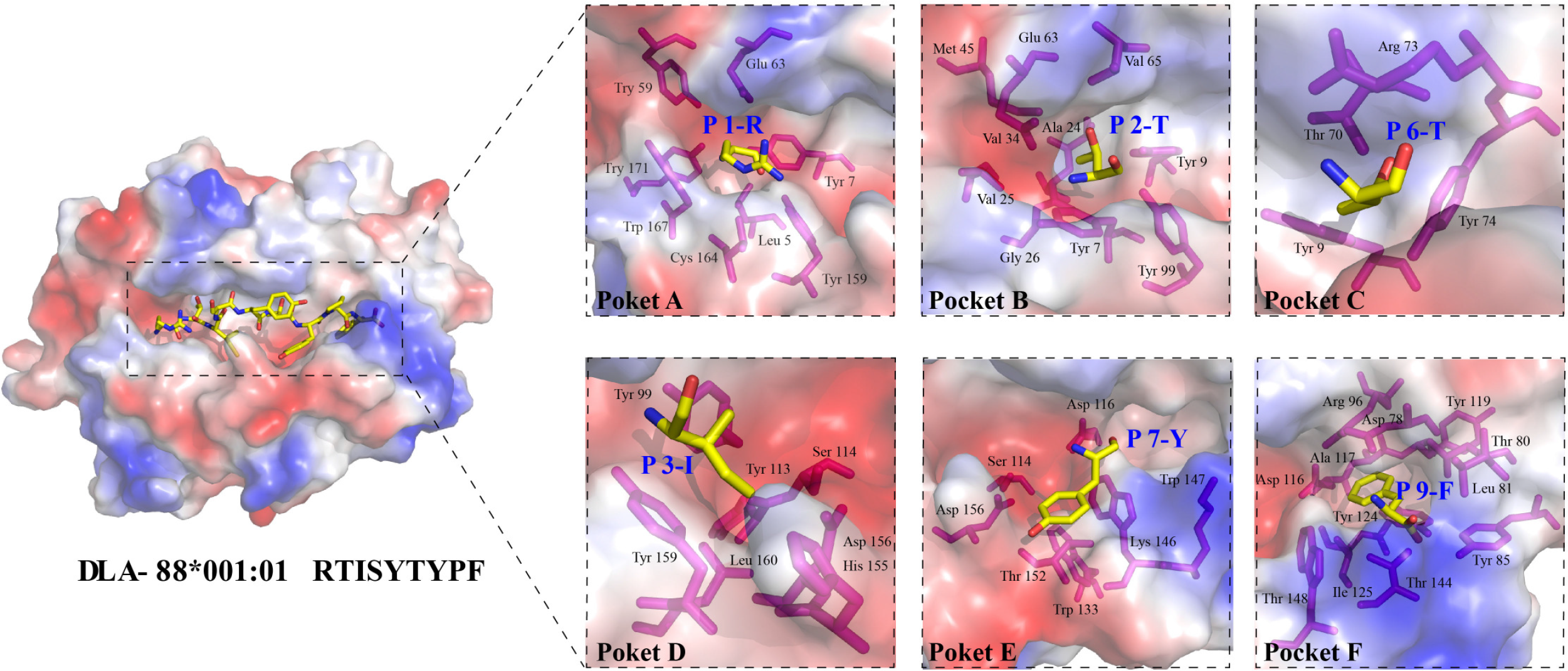
Structural analysis and comparison of six pockets and peptide orientation in the ABG. Composition of the pockets of pDLA-88*001:01. The pockets are shown as surfaces colored by their vacuum electrostatics. Red represents negatively charged residues, blue represents positively charged residues, and gray indicates noncharged residues. The residues composing these pockets are shown as labeled sticks. Residues bound by these pockets are shown as yellow sticks (yellow, carbon; blue, nitrogen; red, oxygen). Pocket A with the P1 residue Arg; Pocket B with the P2 residue Thr; Pocket C with the P6 residue Thr; Pocket D with the P3 residue Ile; Pocket E with the P7 residue Tyr; and Pocket F with the P9 residue Phe.

**Table 2.** The interactions between the peptide and the ABG.

| Hydrogen Bonds and Salt Bridge |  |  |  | Van der Waals' Forces |
| --- | --- | --- | --- | --- |
| Peptide |  | H chain |  |  |
| Residue | Atom | Residue | Atom |  |
| P1-R | O | Y159 | OH | Y59, E63, Y171, Y7, W167, C164, L5, Y159 |
|  | N | Y7 | OH |  |
|  | N | Y171 | OH |  |
| P2-T | N | E63 | OE1 | M45, E63, V65, V34, A24, V25,Y9, G26, Y7, Y99 |
|  | N | E63 | OE2 |  |
|  | OG1 | T66 | OG1 |  |
|  | OG1 | E63 | OE1 |  |
|  | OG1 | E63 | OE2 |  |
| P3-I | N | Y99 | OH | Y99, Y113, S114, Y159, L160, D156, H155 |
| P4-S | O | R73 | NE2 | D156, H155,D156 |
| P6-T | OG1 | R73 | NE | R73, T70, Y9, Y74 |
|  | OE1 | R113 | NH2 |  |
| P8-P | O | W147 | NE1 | D116, W147, K146, T148, I125,Y124 |
| P9-F | OXT | Y85 | OH | R96, D78, Y119, A117, D116, Y124, T80, L81, T148, I125, T144, Y85 |
|  | OXT | T144 | OG1 |  |
|  | N | D78 | OD1 |  |

The structural comparison of pDLA-88*001:01 and pDLA-88*508:01 indicated that these three amino acids are conserved in DLA-88 alleles to form hydrogen bonds with the residue of P1 (Figs. 4, S4A and S4B). In addition, comparisons of the determined pMHC-I molecules in several species, these three amino acids are conserved in all species (Fig. 3).

The B pocket is composed of Met45, Glu63, Val65, Ala24, Val34, Tyr9, Val25, Gly26, Tyr7 and Tyr99. In general, the B pocket is an important anchor site and plays a restrictive role in peptide binding. In both pDLA-88*001:01 and pDLA-88*508:01, the side chains of P2-Thr and P2-Leu are inserted into the B pocket, respectively. In most MHC-I molecules, the residues at positions 63 and 66 are Glu and Lys, respectively, which are both charged residues. In pDLA-88*508:01, the charged Glu63 and Lys66 residues at the top of the B pocket can form two hydrogen bonds with the main chain of P2-Leu, while in pDLA-88*001:01, there is only one hydrogen bond formed with the main chain of P2-Thr, because the residue at position 66 is Val, which is not a charged amino acid. This must affect the restrictive role of the B pocket in pDLA-88*001:01.

The C pocket of pDLA-88*001:01 consists of Arg73, Thr70, Tyr9 and Tyr74 (Fig. 4). The most different C pocket between pDLA-88*001:01 and pDLA-88*508:01 is the residue at position 73. The residue of pDLA-88*001:01 at position 73 is Lys (a charged amino acid), which can form four hydrogen bonds with the peptide CDV-_RTI9_; however, this the residue at position 73 is Val in pDLA-88*508:01 (Fig. S4A and S4B), and this residue creates a restrictive difference in the C pocket of pDLA-88*508:01.

Tyr99, Tyr113, Ser114, Tyr159, Leu160, Asp156 and His155 compose the D pocket of pDLA-88*001:01. The amino acid composition of the D pocket is roughly the same as that of pDLA-88*508:01. However, due to the insertion of Leu155, the D pocket of pDLA-88*508:01 is more stretched, and the buried surface area (BSA) of pDLA-88*508:01 is much larger than that of pDLA-88*001:01. At the same time, because the residue at position 155 of the D pocket of pDLA-88*001:01 is His, which has a large side chain, this side chain points to the ABG, resulting in a smaller D pocket and a stronger selectivity for amino acids to accommodate. The E pocket of pDLA-88*001:01 consists of Lys146, Trp147, Asp116, Trp133, Ser114, Thr152 and Asp156. In most pMHC-I structures, the residue at position 114 is Arg, while in pDLA-88*001:01, it is Ser, which makes the E pocket the largest pocket.

The most important anchor site at the C-terminus of the ABG is the F pocket. Arg95, Ala117, Tyr118, Leu81, Asp77, Thr80, Tyr84, Asp116, Tyr123, Thr143, and Ile124 form the F pocket of pDLA-88*001:01. The side chain of P9-Phe inserts into the F pocket, and there are four hydrogen bonds formed by the surrounding pocket amino acids and main chain of P9-Phe, which may influence peptide binding.

### The different conformations of viral and self-peptide presentations

In most determined pMHC-I molecules, peptides reveal the featured “M” conformation in the ABG. The alignment of CDV-_RTI9_ with other species’ nonapeptides presented by MHC-I shows that CDV-_RTI9_ adopts an overall “M” conformation (Fig. S4C). By superimposing the structures of CDV-_RTI9_ (presented by pDLA-88*001:01) and K9 (presented by pDLA-88*508:01), we found that the side chains of these two peptides are basically the same, and only the side chain of position P7 is significantly different, as shown in Fig. 5. Analysis of the conformation of the self-peptide and viral peptide presented by DLA-88 revealed that the side chains of P1 and P2 of the viral CDV-_RTI9_ peptide are more flexible, while the side chains of P5-P9 are more rigid (Fig. 5A and 5B). However, the stability of the K9 self-peptide is the opposite; the side chains of P1 and P2 are more rigid, while the side chains of P5-P9 are more flexible (Fig. 5C and 5D). The direction difference in the amino acid side chain of the viral peptide and K9 self-peptide is mainly concentrated at the P7 position (Fig. 5E). This difference is mainly caused by the 114th amino acid; the amino acid at position 114 is Ser in DLA-88*001:01 and Arg (which has a large side chain) in DLA-88*508:01. The small side chain of Ser can accommodate the large side chain of CDV-_RTI9_ P7-Tyr, while the large side chain of Arg occupies the position of the K9 self-peptide P7-Leu.

**Fig. 5.**
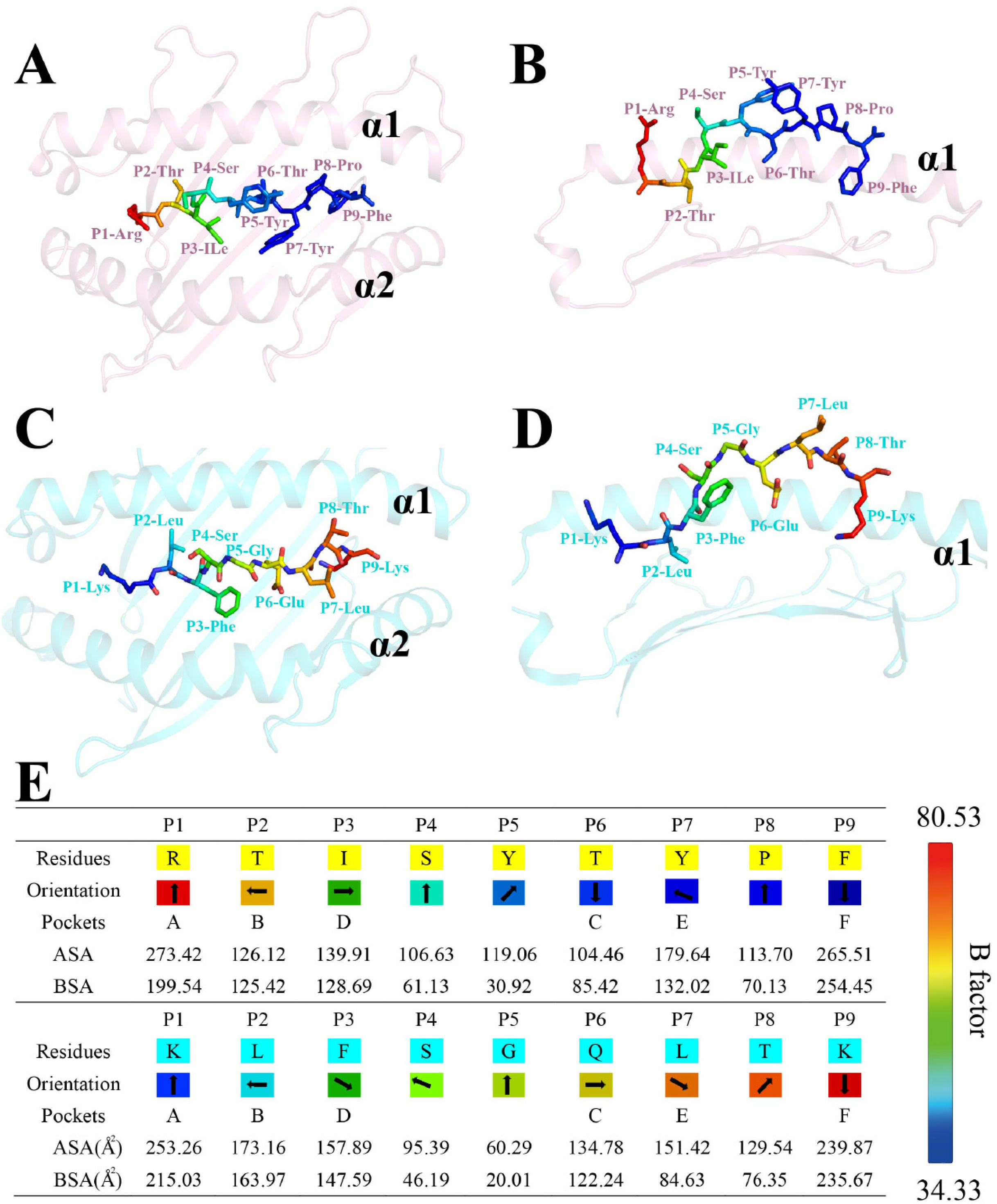
Structure of the peptides presented by pDLA-88*001:01 and pDLA-88*50801. (**A**) CDV-H1_RF9_ presented by pDLA-88*001:01. Peptides are shown as cartoon models and colored based on isotropic B factors. (**B**) View of CDV-H1_RF9_ from the α2 helices. (**C**) K9 self-peptide presented by pDLA-88*50801. (**D**) View of K9 from the α2 helices. (**E**) Side chain orientation of peptides in the pDLA-88*001:01 and pDLA-88*50801 structures in the side view from the peptide N-terminus to the C-terminus, as viewed in profile from the peptide N-terminus toward the C-terminus. An arrow pointing up indicates that an amino acid residue is oriented toward the TCR, down is toward the floor of the ABG, left is toward the α1 helix domain, and right is for the α2 helix domain. The accessible surface area (ASA) and BSA represent the exposed and buried surface areas of each peptide residue, respectively.

To confirm the restrictive pockets and the primary anchor residues of pDLA-88*001:01, alanine mutation and circular dichroism (25) spectroscopy experiments were used to investigate CDV-_RTI9_ (Fig. 6). The wild-type CDV-_RTI9_ peptide was used as a control, and the results showed that the midpoint transition temperature (Tm) peaks of the pDLA-88*001:01 complex with P2-Ala, P2-Gly and P9-Ala peptides were significantly decreased (44°C, 46°C and 44°C, respectively, Fig. 6). The Tm value of the wild-type peptide was 50°C. Based on structural and refolding data, we summarized the primary peptide pDLA-88*001:01 binding motif as follows: X-(P/T/A/V/L/S)-X-X-X-(E/F/L/V/A/G/Q/S/Y/T)-X-X-X-X-(F/L/R/K/I).

**Fig. 6.**
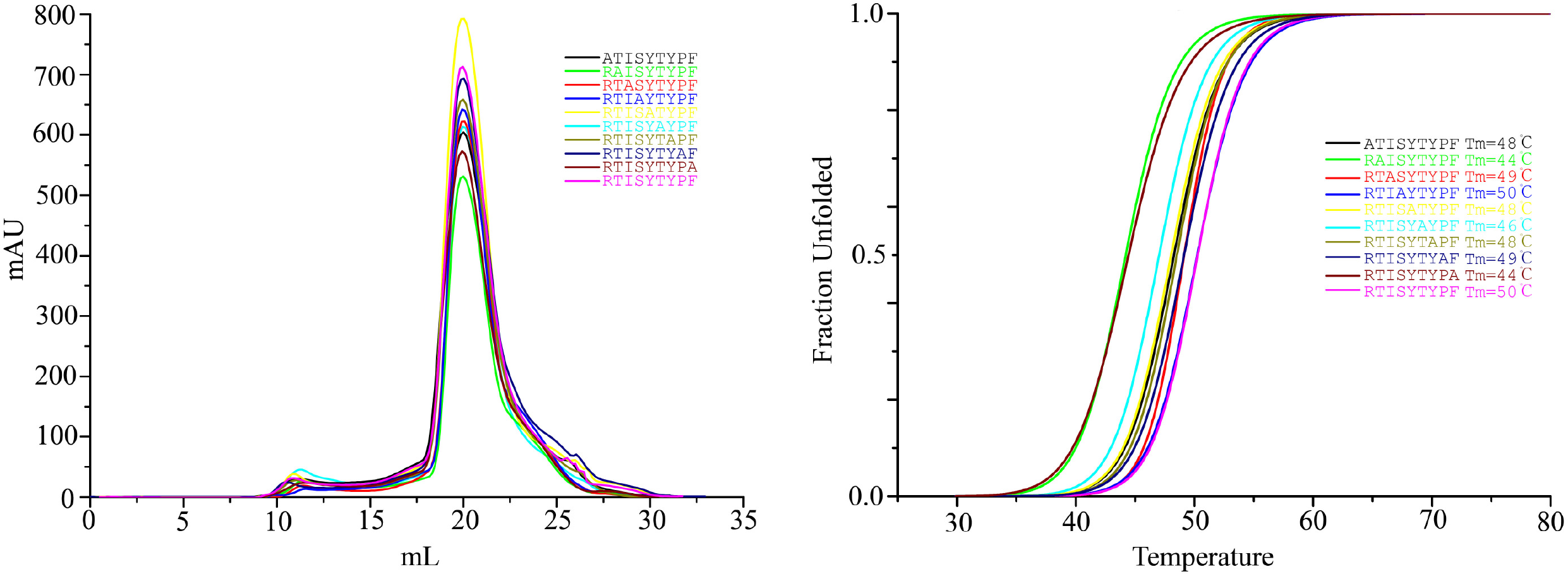
Peptide-induced assembly and DLA-88*001:01-CDV-H1_RF9_ complex stabilization were tested using *in vitro* refolding. (A) Gel filtration chromatograms of the refolded products. The main peaks represent the correctly refolded DLA-88*50801-CDV-H1_RF9_ complex (35 kDa). The refolding efficiencies are represented by the relevant concentration ratio and the height of the DLA-88*001:01-CDV-H1_RF9_ complex peak for each mutant. A high DLA-88*001:01-CDV-H1 RF9 complex peak indicated the superior ability of the peptide to promote MHC renaturation. Otherwise, if a small or absent DLA-88*001:01-CDV-_RTI9_ complex peak was observed, the peptide was not considered to stabilize the complex and was therefore treated as a non-presented peptide. The mutated peptide P2-A clearly yielded the lowest refolding efficiency. (**B**) Thermal stabilities of the pDLA-88*001:01 complex. DLA-88*001:01 bound to CDV-H1_RF9_ or one of 9 mutant peptides was tested by CD spectroscopy. The denaturation curves of the complexes with the different peptides are indicated in different colors.

### Selective method for TCR recognition of DLA-88*001:01

In a previous study by Liu Jun (2016), the DLA-88*508:01 H chain with a Leu155 insertion, causing a change in the height of the position interacting with the TCR. As shown in Fig. 7A, pDLA-88*508:01 Leu155 is 2.9 Å higher than pDLA-88*001:01 and 3.3 Å higher than human leukocyte antigen-A2 (HLA-A2, PDB ID: 3OXS). All of the residues on the α1 and α2 helices of DLA-88 that contact TCRs were analyzed (Fig. 7B and 7C). DLA-88*508:01 has more amino acid residues than pDLA-88*001:01 and can interact with TCRs (Fig. 7D). Therefore, pDLA-88*508:01 is quite different from other known mammalian MHC-I structures, which Liu et al speculated may be a novel pattern of MHC-I recognition of the TCR. In the pDLA-88*001:01 we reported, the H chain did not have this Leu155 insertion, and our structure is similar to the most known pMHC-I structures. Because the ABG of pDLA-88*001:01 has one amino acid less than pDLA-88*508:01, the highest interaction region between ABG and TCR is lower than that of pDLA-88*508:01, and it also affects the interaction between DLA H and light (L) chains. As mentioned above, the interaction between the H and L chains of DLA-88*001:01 has more hydrogen bonds and amino acids than pDLA-88*508:01. The interaction area between the H and L chains of pDLA-88*001:01 is also different from that of pDLA-88*508:01. As shown in Fig. 8, the area of DLA-88*001:01 buried in the L chain is 1333.65 Å^2^ for pDLA-88*001:01-CDV-_RTI9_, while the are of DLA-88*508:01 buried in the L chain is 1183.24 Å^2^. Similarly, the area of the L chain buried in DLA-88*001:01 is also larger than that of pDLA-88*508:01. Therefore, we speculated that there may be two modes by which pMHC-I recognizes TCRs in canine DLA-88.

**Fig. 7.**
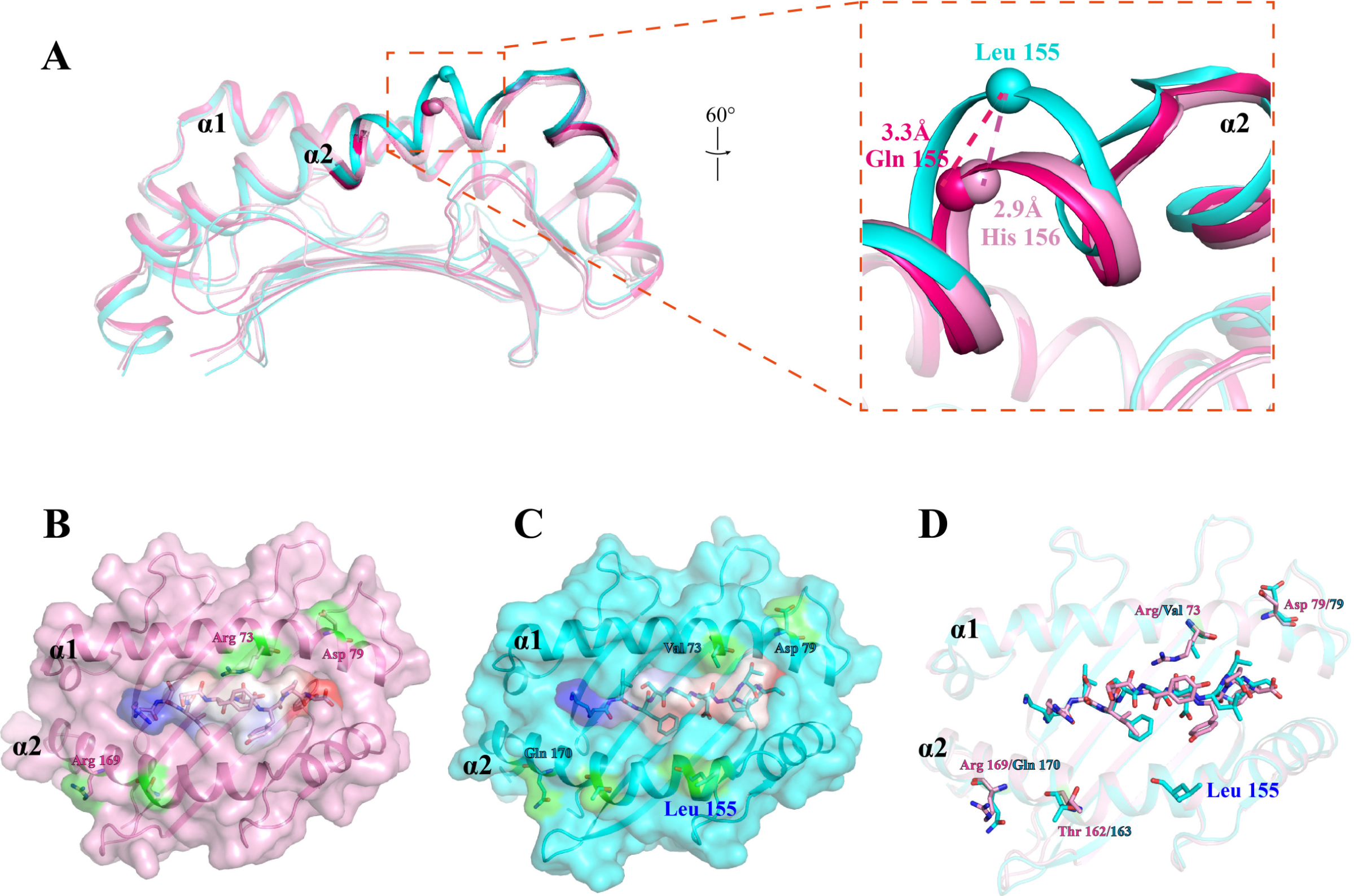
(A) The ABG differences in DAL-88*001:01 and DAL-88*50801. DAL-88*001:01 is shown in cyan cartoon form, DAL-88*001:01 is shown in pink cartoon form, and HLA-A2 (PDB ID: 3OXS) is shown in magenta cartoon form. The highest residue of the α2 helix in the ABG of DAL-88*50801 is shown as a sphere, and the corresponding positions of DAL-88*001:01 and HLA-A2 are shown as pink and magenta spheres, respectively. (**B**) Solvent-exposed residues (green surface) on the α1 and α2 helices of DLA-88*001:01 that may contact TCRs. (**C**) Solvent-exposed residues (green surface) on the α1 and α2 helices of DLA-88*50801 that may contact TCRs. (**D**) Superimposition of the ABG of pDLA-88*001:01 and of pDLA-88*50801; residues that may contact TCRs, are shown in pink and cyan sticks, respectively.

**Fig. 8.**
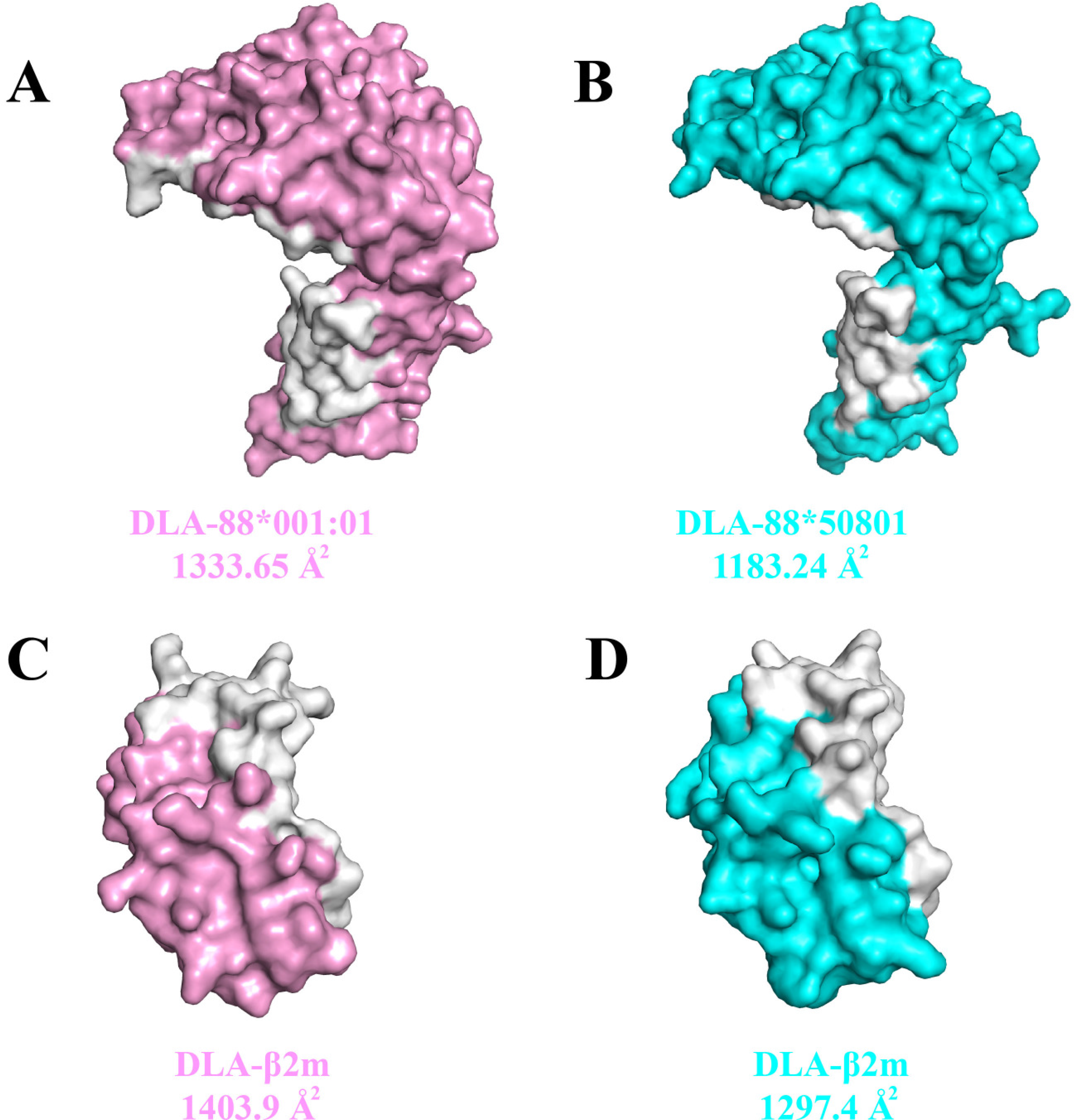
Interface areas of the H and L chains of pDLA-88*001:01 and pDLA-88*50801. (**A**) The H chain of pDLA-88*001:01 is shown in pink surface form. The white areas indicate the interactive regions contacting the L chain. Detailed interface areas are shown under the structure. (**B**) The H chain of pDLA-88*50801 is shown in cyan surface form. The white areas indicate the interactive regions contacting the L chain. (**C**) The L chain of pDLA-88*001:01 is shown in pink surface form. The white areas indicate the interactive regions contacting the H chain. Detailed interface areas are shown under the structure. (**D**) The L chain of pDLA-88*50801 is shown in cyan surface form. The white areas indicate the interactive regions contacting the H chain.

### Viral peptides matched the motif of pDLA-88*001:01

Using the motif of pDLA-88*001:01 that we identified, the proteomes of Influenza A virus subtype H7N9, CPV, canine measles virus (CMV) and CDV strains were screened to identify the epitope peptides that could be presented by pDLA-88*001:01 (Fig. 9). Three hundred and twenty-four peptides matching the peptide-binding motif of pDLA-88*001:01 were determined (Table S2). We selected five nonapeptides from these peptides to test their binding with DLA-88*001:01 by *in vitro* refolding (Table S3). The results show that the predicted epitopes could bind to DLA-88*001:01. In summary, we predicted twenty-three peptides from the neuraminidase protein of H7N9, thirty-one nonapeptides from the hemagglutinin protein (H protein) of H7N9, fourteen peptides from the VP2 capsid protein of CPV, seven peptides from the VP1 capsid protein of CPV, thirty-five peptides from the nuclear phosphoprotein (NP) of CPV, nine peptides from the phosphoprotein (P protein) of CMV, thirty-six peptides from NP of CMV, fifty-nine peptides from the H protein of CMV, twenty-seven nonapeptides from the P protein of CDV, thirty-two nonapeptides from the N protein of CDV, and fifty-one nonapeptides from the H protein of CDV that have high affinity and have the binding motif of DLA-88*001:01. This study has important significance for the design of peptide vaccines.

**Fig. 9.**
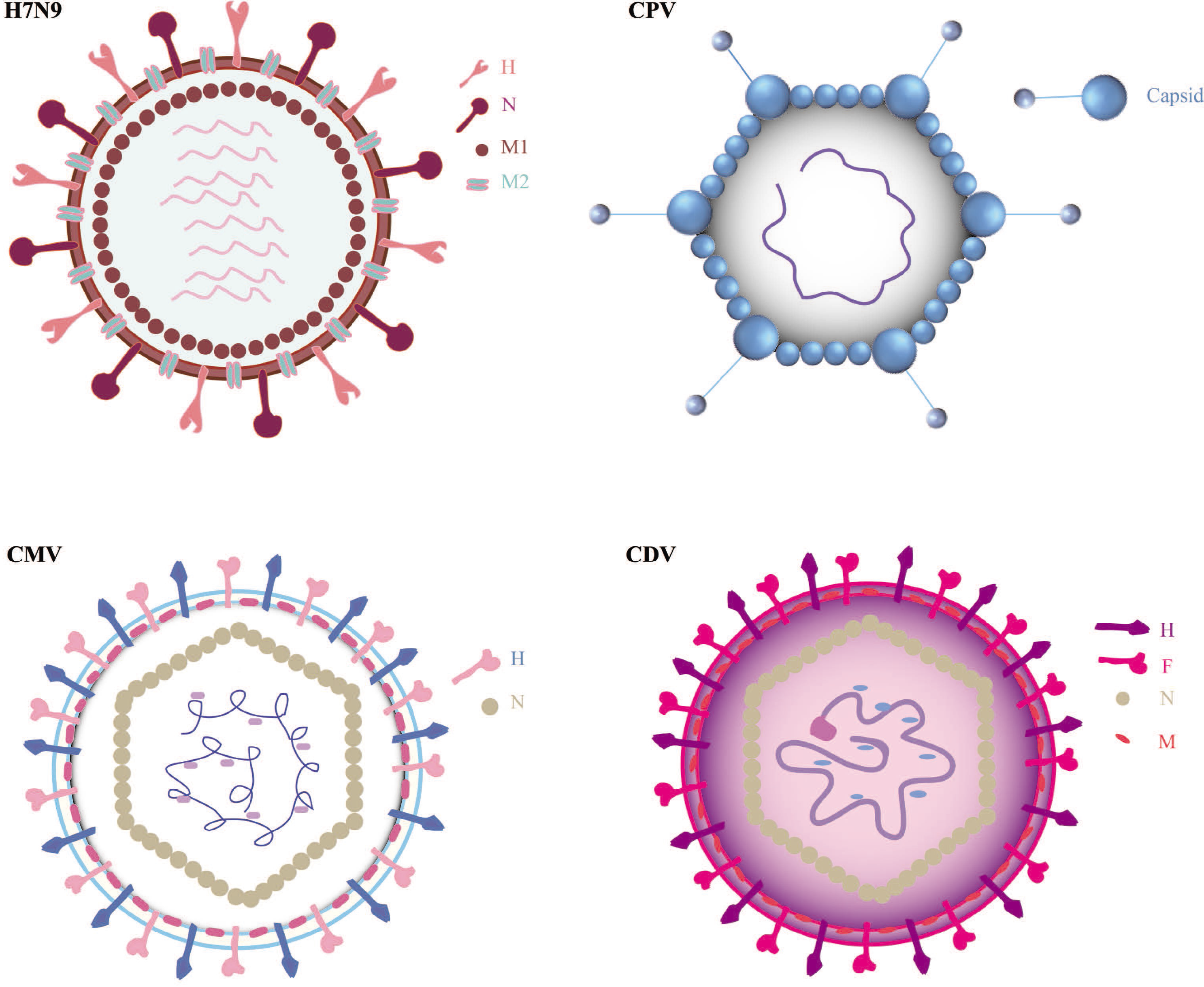
Peptide predictions from different viruses according to the binding motifs of pDLA-88*001:01.

## Discussion

Since there is only one classical MHC-I locus and 76 alleles in dogs, it is easier to study antiviral CTL immunity because the determination of the polymorphism of DLA-88 alleles is accessible (26). In this paper, the structure of the normal-length DLA-88 and pDLA-88*001:01 complex was determined for the first time, and the characteristics of pDLA-88*001:01 preferentially presenting viral peptides were elucidated. In addition, the structural differences between DLA-88*508:01 and pDLA-88*001:01 were analyzed. According to the features of pDLA-88*001:01, a variety of canine viral epitopes were found, which provided CTL candidate epitopes for multi-epitope vaccines.

pDLA-88*001:01 showed classical pMHC-I characteristics in the α1, α2, α3 and β2m domains. CDV-_RTI9_ is a typical “M” conformation in the ABG that is thought to activate T cells (27, 28). The results of amino acid sequence and 3D structure comparisons showed that DLA-88*001:01 was similar to most of the pMHC-I topologies but was significantly different from the reported pDLA-88*508:01 3D structure. There was a Leu insertion in the α2 region of DLA-88*508:01. The interaction region between the ABG and TCR was approximately 2.9 Å (22) higher in DLA-88*508:01 than in pDLA-88*001:01. There were differences in amino acid sequences and lengths between species or within species of classical MHC-I molecules. These differences lead to differences in MHC-Ia polymorphisms, ABG micro polymorphisms, and crosses and overlaps of antigen presentation profiles (25). This is the strategy of MHC-I system evolution. Therefore, it is not uncommon to have amino acid insertions or deletions in the ABG region. For example, the amphibian *Xenopus*, whose α2 domain helix kink region of the ABG has two amino acid EV insertions, results in the highest point of the ABG being approximately 3.8 Å higher than that of other species, and all frogs have the insertion of these two amino acids, making this frog-specific characteristic (29). There are some species-specific characteristics with the insertion of N-terminal residues in the ABG region of bats (30). Amino acid insertion in mammalian MHC-Ia occurs in most species, and the length of the ABG in the same species is rarely changed. DLA-88*001:01 and DLA-88*508:01 are both alleles at the DLA-88 locus in dogs. The difference in amino acids between DLA-88*001:01 and DLA-88*508:01 changes the motif of the ABG-binding peptide and antigen-binding profile and affects the interaction mode with the canine TCR. This is also the evolutionary feature of the single classic DLA-88 in canines; although there is only one classical MHC-I locus, which expresses two kinds of alleles, the insertion or deletion of amino acids can change the ABG polymorphism and complete a high-throughput specific cellular immune response against various viral pathogens.

DLA-88*001:01 binds to the CDV peptide, which is different from that of DLA-88*508:01 (26). Canine distemper is a global epidemic. Effective measures to prevent canine distemper mainly rely on attenuated vaccines or oil emulsion-inactivated vaccines (31). In this study, the complex structure of pDLA-88*001:01 and the CDV peptide was solved using X-ray crystallography. Through the structural characteristics of the CDV-_RTI9_ alanine mutation, the CDV peptide binding rule of DLA-88*001:01 was found, and the binding peptide motif of pDLA-88*001:01 was obtained. According to the binding peptide motif, the whole protein sequence of CDV, H7N9, CPV, CMV and other viral strains in canines was scanned, and the epitope peptide profile of DLA-88*001:01 was determined. Finally, 324 peptides were obtained that matched the binding motif of pDLA-88*001:01 (Table S2). Among them, there were 23 peptides from the N protein of H7N9, 31 from the H protein of H7N9, 14 from VP2 of CPV, 7 from VP1 of CPV, 35 from NP of CPV, 9 from the P protein of CMV, 36 from NP of CMV, 59 from the H protein of CMV, 27 nonapeptides from the P protein of CDV, 32 nonapeptides from the N protein of CDV and 51 nonapeptides from the H protein of CDV. Although this series of candidate CTL epitopes needs to be screened, the research scope of CTL epitopes of the dog DLA-88 virus is provided.

In conclusion, the pDLA-88*001:01 reported in this paper shows the difference in the 3D structure between another class of multiple DLA-88 alleles in canines, and this study is the first to describe the characteristics of ABG binding to the CDV peptide, and rationally infers more than 300 candidate CTL epitopes of various canine viruses. These results promote the research of antiviral cellular immunology and multi-epitope vaccines in dogs.

## Materials and methods

### Gene structure and sequence analyses of MHC-I alleles in dogs

According to Immuno-Polymorphism Database (IPD) (32) (https://www.ebi.ac.uk/ipd/mhc/), National Center for Biotechnology Information (NCBI) (https://pubmed.ncbi.nlm.nih.gov) and reported canine MHC-I/DLA literature, DLA has four loci located on chromosomes 18 and 12. The canine MHC-I loci are named DLA-88, DLA-12, DLA-64, and DLA-79 (18). In 2000, the DLA Nomenclature Committee of the International Society for Animal Genetics (ISAG) determined that DLA-88 was the only classical allele, and the other three loci were nonclassical alleles (33). There are 76 alleles in the DLA-88 locus; thus, we collected the extracellular domain of DLA-88 alleles, namely, the α1 and α2 regions, and analyzed the gene sequence and evolution. Multiple classical *DLA-88* genes were aligned using the ClustalW Sequence Alignment program of the comprehensive suite of molecular biology and sequence analysis tools in Geneious Prime. The phylogenetic tree, consisting of 76 *DLA-88* genes determined in this study, was constructed with only synonymous substitutions that were used to identify differences by the neighbor-joining (NJ) (34) method in Geneious Prime. Evolutionary genetic distance nucleotides were estimated by the Jukes-Cantor method using Geneious Prime. The tree was edited in another online software (Evolgenius; https://evolgenius.info/evolview-v2).

### Synthesis of viral peptides

A total of 14 viral nonapeptides were used in these experiments (Tables 1 and 2). These nonapeptides matched the sequences of the CDV, rabies virus and influenza virus that were predicted by NetMHCpan-4.0 (http://www.cbs.dtu.dk/services/NetMHCpan). The CA-7 peptide is a self-peptide that has been reported (22). These peptides were synthesized and purified by reversed-phase high-performance liquid chromatography (SciLight Biotechnology, Beijing, China) with >95% purity. These lyophilized peptides were stored at −20°C, were dissolved in DMSO at a concentration of 25 mg/mL before use and then used as described.

### Protein preparation

The gene fragments encoding the extracellular domain (1-275 amino acids) of the DLA-88*001:01 allele (GenBank accession No. JQ733514) and extracellular domain (1-98 amino acids) of canine β2m (GenBank accession No. JQ733515) were synthesized by Shanghai Invitrogen Life Technologies. The DLA-88*001:01 and canine β2m genes were digested with the *Nde*I and *Xho*I enzymes, ligated into the pET21a expression vector (Novagen, Merck KGaA, Darmstadt, Germany) and then transformed into *Escherichia coli* BL21(DE3) cells. Recombinant DLA-88*001:01 was expressed in inclusion bodies and purified as described previously (29). Recombinant canine β2m was also expressed in inclusion bodies and purified as previously described (7). Finally, the inclusion bodies of DLA-88*001:01 and canine β2m were dissolved in 6 M guanidine hydrochloride solution (6 M guanidine hydrochloride (Gua-HCl), 50 mM Tris-HCl (pH 8.0), 10 mM EDTA, 100 mM NaCl, 10% v/v glycerol and 10 mM DTT). The final concentration was 30 mg/ml. The proteins were stored at −20°C.

### The assembly and purification of the DLA-88*001:01 complex

Refolding of DLA-88*001:01 with the CDV-H1_RF9_ viral peptide. To form a complex with each peptide (Table 2), DLA-88*001:01, β2m and the nonapeptide were refolded at a ratio of 3:1:1 using the gradual dilution method, as previously described (35). As a negative control, DLA-88*001:01 and β2m were also refolded without the peptide. After 48 h of incubation at 4°C, the remaining soluble portion of the complex was concentrated and then purified by chromatography on a Superdex 200 16/60 column followed by Resource Q anion-exchange chromatography (GE Healthcare), as previously described (36). Finally, the purified complex protein solution was replaced with 10 mM Tris-HCl and 50 mM NaCl (pH 8.0) three times.

### Thermostability measurements using circular dichroism spectroscopy

The thermostabilities of DLA-88*001:01 with nonapeptides were tested by CD spectroscopy. CD spectra were measured at 20°C on a Jasco J-810 spectropolarimeter equipped with a temperature-controlled cell holder. The protein concentration was 0.1 mg/mL in Tris buffer (20 mM Tris and 50 mM NaCl (pH 8.0)). Thermal denaturation curves were determined by monitoring the CD value at 218 nm using a 1-mm optical-path-length cell as the temperature was raised from 25 to 80°C at a rate of 1°C/min. The temperature of the sample solution was directly measured with a thermistor. The fraction of unfolded protein was calculated from the mean residue ellipticity (θ) using a standard method. The unfolded fraction (%) is expressed as (Θ-ΘN)/(ΘU-ΘN), where ΘN and ΘU are the mean residue ellipticity values in the fully folded and fully unfolded states, respectively. The Tm was determined by fitting the data to the denaturation curves using the Origin 8.0 program (OriginLab) as described previously (37). Based on the CD spectroscopy results, the nonapeptide-binding motifs of DLA-88*001:01 were further extrapolated from the potential CTL epitopes of viruses detected from dogs.

### Crystallization and data collection

One viral peptide, CDV-H1_RF9_ (RTISYTYPF, residues 543-551), was selected for crystallization with the DLA-88*001:01 H chain and canine β2m. The DLA-88*001:01-CDV-H1_RF9_ complex was concentrated to 6 and 12 mg/mL in buffer containing 20 mM Tris (pH 8.0) and 50 mM NaCl for crystallization. After being mixed with reservoir buffer at a 1:1 ratio, the purified DLA-88*001:01-β2m-peptide complex (pDLA-88*001:01) was crystallized by the hanging-drop vapor diffusion method at 291 K. Polyethylene glycol (PET)/ion kits (Hampton Research, Riverside, CA) were used to screen for crystals. After several days, crystals of the DLA-88*001:01-CDV-H1_RF9_ complex were obtained with solution 33 (20% (*w/v*) PEG 3350 and 0.2 M sodium sulfate decahydrate) at a concentration of 12 mg/mL. Diffraction data were collected using an in-house X-ray source (Rigaku MicroMax007 desktop rotating anode X-ray generator with a Cu target operated at 40 kV and 30 mA) and an R-Axis IV^++^ imaging plate detector at a wavelength of 1.5418 Å. In each case, the crystal was first soaked in a reservoir solution containing 15% glycerol as a cryoprotectant for several seconds and then flash-cooled in a stream of gaseous nitrogen at 100 K (12). The collected intensities were indexed, integrated, corrected for absorption, scaled, and merged using HKL2000 (38).

### Structure determination and refinement

The structure of the DLA-88*001:01-CDV-H1 _RF9_ complex was solved by molecular replacement using the MOLREP program with SLA-1*0401 (PDB ID: 3QQ3) as the search model. Extensive model building was performed by hand using Coot (39), and restrained refinement was performed using REFMAC5. Further rounds of refinement were performed using the phenix.refine program implemented in the PHENIX package with isotropic ADP refinement and bulk solvent modeling, which improved the R and R_free_ factors from 0.194 and 0.209 to 0.151 and 0.177, respectively. The stereochemical quality of the final model was assessed with the PROCHECK program (40). Data collection and refinement statistics are listed in Table 1.

### Structure determination, refinement and data analysis

Analysis of the determined DLA-88*001:01-CDV-H1_RF9_ structure was performed using PyMOL (https://www.pymol.org) (Schrodinger, 2010), Collaborative Computational Project Number 4 (CCP4) (41) and Coot. The SignalP 3.0 server was used to predict the presence and location of signal peptide cleavage sites (42). The comparison of amino acid sequences from different proteins was performed using ClustalW2 (http://www.ebi.ac.uk/Tools/msa/clustalw2/) (43).

### Protein structure accession numbers

The coordinates and structure factors of the DLA-88*001:01-CDV-H1 RF9 complex have been deposited in the Protein Data Bank with the accession number 7CJQ.

## Acknowledgments

This work was supported by the National Natural Science Foundation of China (Grant NO. 31972683 and Grant NO. 31572493).

C.X. designed the project; YS., L.M., S.L., YW., Q.Y and Q.J. performed the experimental work; Y.S. prepared the proteins; Y.S, L.M., S.L. and Y.W. performed the CD spectra analysis and crystallization experiments; S.L. solved and uploaded the structure to the PDB bank; S.L., L.M. and Y.W. analyzed the original data; the draft of the manuscript was written by L.M. and S.L.; revisions were made by L.M., S.L. and C.X.

We acknowledge the assistance of the staff of the Shanghai Synchrotron Radiation Facility of China.

## Supplementary Material

**Fig. S1. The amino acid sequence alignment of the DLA-88 alleles**. The amino sequences of all published DLA-88 alleles are presented in the clustered alignment and were generated by ClustalW. Variations between the alleles in the polymorphic α2 domain are indicated. The insertion of the 155^th^ amino acid is framed in black.

**Fig. S2. Superposition of pDLA-88*001:01 and pDLA-88*50801.** pDLA-88*001:01 is shown in pink cartoon form, and pDLA-88*50801 is shown in cyan cartoon form. The RMSD values of the ABG, β2m and α3 domain are labeled.

**Fig. S3. Interactions between the H and L chains of pDLA-88*50801.** The H chain is shown in cyan, with the interactive residues shown as sticks and colored according to the atom type (blue, nitrogen; red, oxygen). The interactive residues on the L chain are colored in blue. The hydrogen bonds are shown as yellow dashed lines. All of the interactions and interactive residues are magnified within a corresponding color dotted box and labeled.

**Fig. S4. **(A)** Interactions of CDV-H1_RF9_ and the ABG of pDLA-88*001:01.** The residues in the ABG that can form hydrogen bonds with CDV-H1_RF9_ are shown as sticks. The hydrogen bonds are shown as yellow dashed lines. The different residues with pDLA-88*50801 are labeled in pink. (**B**) Interactions of K9 and the ABG of pDLA-88*50801. The residues in the ABG that can form hydrogen bonds with K9 are shown as sticks. The different residues with pDLA-88*001:01 are labeled in cyan. (**C**) Structural alignment of the peptide conformation of DLA-88 and MHC-I of other species. Characteristics of the peptide conformation of DLA-88, which is shown as ribbons with a thick line in hot pink. The remaining peptides are shown in different colors depending on their species and are depicted as ribbons.

**Table S1. Predicted peptides and their binding to DLA-88*001:01/β2m evaluated by *in vitro* refolding.**

**Table S2. Predicted peptides that can bind to DLA-88*001:01 β2m.**

**Table S3. Selected peptides and their binding to DLA-88*001:01/β2m verified by *in vitro* refolding.**

